# Data-optimal scaling of paired antibody language models

**DOI:** 10.1101/2025.09.02.673765

**Authors:** Mahdi Shafiei Neyestanak, Sarah M. Burbach, Karenna Ng, Praneeth Gangavarapu, Jonathan Hurtado, Judie Magura, Nasreen Ismail, Daniel Muema, Thumbi Ndung’u, Andrew B. Ward, Bryan Briney

## Abstract

Scaling laws for large language models in natural language domains are typically derived under the assumption that performance is primarily compute-constrained. In contrast, antibody language models (AbLMs) trained on paired sequences are primarily data-limited, thus requiring different considerations. To explore how model size and data scale affect AbLM performance, we trained 15 AbLMs across all pairwise combinations of five model sizes and three training data sizes. From these experiments, we derive an AbLM-specific scaling law and estimate that training a data-optimal AbLM equivalent of the highly performant 650M-parameter ESM-2 protein language model would require ∼5.5 million paired antibody sequences. Evaluation on multiple downstream classification tasks revealed that significant performance gains emerged only with sufficiently large model size, suggesting that in data-limited domains, improved performance depends jointly on both model scale and data volume.

## INTRODUCTION

Extracting the structural and functional information stored in protein sequences is a long-standing and fundamental biological problem (1). Language models (LMs), originally developed for natural language processing (NLP), have been broadly adapted to biological sequences with the goal of better understanding the “language” of proteins and antibodies. Transformer-based (2) protein language models (pLMs) such as ProteinBERT (3), the ProtTrans model family (4), and the Evolutionary Scale Modeling (ESM) series (5–7) have emerged as a transformative paradigm for learning context-aware representations of amino acid sequences with a variety of biological and clinical applications (8–10).

Antibodies are highly diverse, with previous studies estimating that the circulating antibody repertoire contains as many as 10^18^ unique paired antibodies (11). Prior to antigen exposure, antibody repertoire diversity is achieved through the recombination of modular variable (V), diversity (D), and joining (J) germline gene segments. The majority of pre-immune repertoire diversity is concentrated in the complementarity-determining regions (CDRs) of antibody heavy and light chains, the result of non-templated addition at the junctions between recombined germline gene segments. Upon antigen recognition, antibodies are affinity matured by somatic hypermutation, which introduces mutations into the B cell receptor (BCR) sequence, followed by antigen-driven selection of productive mutations (12). Characterization of these functional antibody profiles elucidates critical mechanisms underlying both protective immunity and immunopathogenesis (13–16) and even proposes potential therapeutic solutions (17–19).

Previous studies have shown that specialized antibody LMs (AbLMs), as opposed to repurposed pLMs or other general biological LMs, are more useful for antibody-specific tasks (20–22). However, the modular nature of antibody recombination results in large regions of tokens that are well conserved even among affinity-matured antibodies. This means that most tokens in an antibody sequence have relatively little training value and may even inhibit the model from learning useful features like somatic hypermutation (21). Focusing training on non-templated heavy and light chain CDR3s using techniques such as preferential masking (23) or focal loss (21,24) can improve model performance, mediated by a better understanding of these complex and information-dense regions.

In addition, antibodies are composed of a unique pairing of heavy and light chains, and cross-chain structural and functional features are critical to antibody specificity and antigen binding kinetics (25,26). We and others have shown that training AbLMs using natively paired antibody sequences results in improved model performance (21,22,27), despite the relative paucity of paired antibody sequence datasets (28,29). Moreover, our recent study revealed that incorporating unpaired sequences makes it possible to train larger models, but yields only marginal gains in downstream task performance compared to training with exclusively paired sequences (30). This suggests that optimizing the training of exclusively paired AbLMs, which has thus far been overlooked, is an essential next step.

Optimal LM training involves carefully balancing model size, training data scale, and compute resources (31). Here again, AbLM training requires a different set of considerations than pLMs or NLP LMs. Within reasonable limits, transformer-based LMs generally improve as their parameter count increases (32,33). NLP LMs are principally compute-constrained, so much work on NLP LM scaling has focused on discovering compute-optimal scaling laws (31,34).

Recent work on pLM scaling has similarly focused on compute-optimality (35,36). In contrast, training paired AbLMs is primarily data-constrained, a fact that is exacerbated by the low training value of many tokens in an antibody sequence. Here, we address the question: Given a fixed number of training examples and unconstrained compute, what is the optimal AbLM model size? We systematically pretrained AbLMs across five different model sizes and three training data scales. These models facilitate a rigorous evaluation of performance scaling trends to define the rules for data-optimal training of natively paired AbLMs.

## RESULTS

### Models and training data

We trained a series of AbLMs of five different sizes: 8, 35, 150, 350, and 650 million parameters (***Figure 1***). All models used an ESM-2 architecture (6), which includes rotary positional embeddings (37) and pre-layer normalization (38). All models were trained using an identical training schedule. To ensure fair model comparisons, checkpoint selection was guided by performance on the held-out evaluation dataset. The optimal checkpoint was defined as the point at which evaluation loss began diverging from training loss, indicating the onset of overfitting. The chosen checkpoint for each model is provided in ***Table S1***.

**Figure 1.**
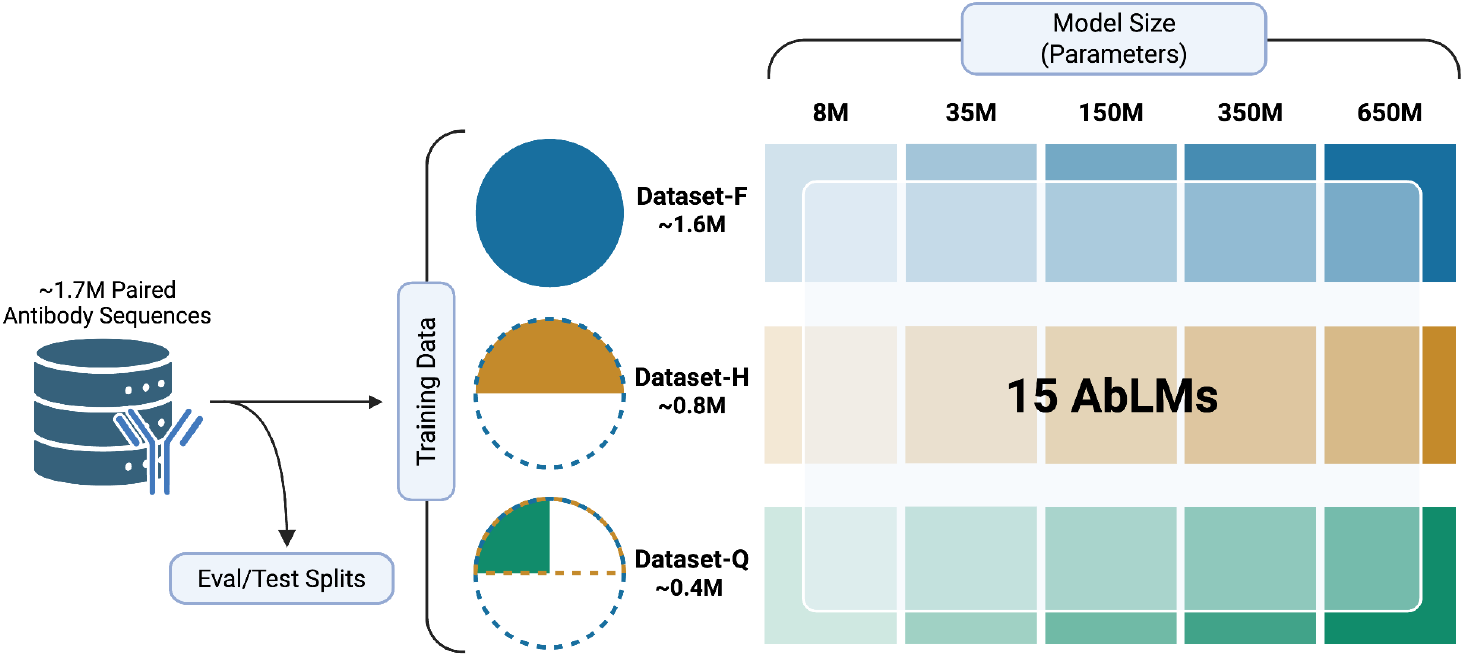
Experimental design for evaluating AbLM scaling dynamics. Fifteen models were pretrained with combinations of five model sizes (8M, 35M, 150M, 350M, and 650M parameters) and three training-data scales (Dataset-F: 1.6 M sequences; Dataset-H: 0.8 M; Dataset-Q: 0.4 M). Separate evaluation and test sets were held out from the training data. Created with BioRender.com.

We constructed our dataset from antibody repertoires of healthy donors, applied quality filtering (see Methods), and clustered at 90% sequence identity to minimize redundancy. To ensure consistency across experiments, fixed evaluation (2%) and test (2%) subsets were extracted from the full dataset before any training subset sampling. The remaining 96% of the paired antibody sequences were used to generate nested structure training sets. AbLMs of each size were trained on three datasets: the full training set of paired antibody sequences with ∼1.6M sequences (Dataset-F), half of the dataset with ∼800k sequences (Dataset-H), and a quarter of the dataset with ∼400k sequences (Dataset-Q). Training datasets were randomly sampled from the next largest dataset to give us a nested structure, allowing us to isolate the effects of model capacity and training data volume. Pairwise χ^2^ tests of independence on V, J, and V/J gene usage composition showed no significant differences between the three subsampled training datasets (***Figure S1, Table S2***). For clarity and brevity, we will denote model configurations as *{model size}-{training data scale}* (e.g., 35M-Q), where “F,” “H,” and “Q” represent Dataset-F, Dataset-H, and Dataset-Q, respectively.

### Identifying data-optimal AbLM sizes using FixedData profiles

Inspired by the IsoFLOP profiles previously used to assess compute optimality (31), we developed “FixedData profiles” by measuring the performance of differently sized models while keeping the training data scale constant. Each of the 15 pretrained models (***Figure 1***) was evaluated using a masked language modeling (MLM) objective across 10 independent evaluation datasets of paired antibody sequences, each derived from distinct donors not present in the training, evaluation, or test datasets. A quadratic regression was fit to the average cross-entropy loss across all test datasets for each model ***(Figure 2a, Table S3***), with outlier points far from the loss minimum excluded from the fit. The complete set of points, including those excluded from the fit, is shown in ***Figure S2***. As expected, models of the same size showed improved performance with increasing training data scale. The optimal model size, identified as the minimum of the FixedData curve, also increased with training data scale. We then fit a power law to the optimal model sizes to extrapolate the optimal amount of training data for any model size (***Figure 2b***).

**Figure 2.**
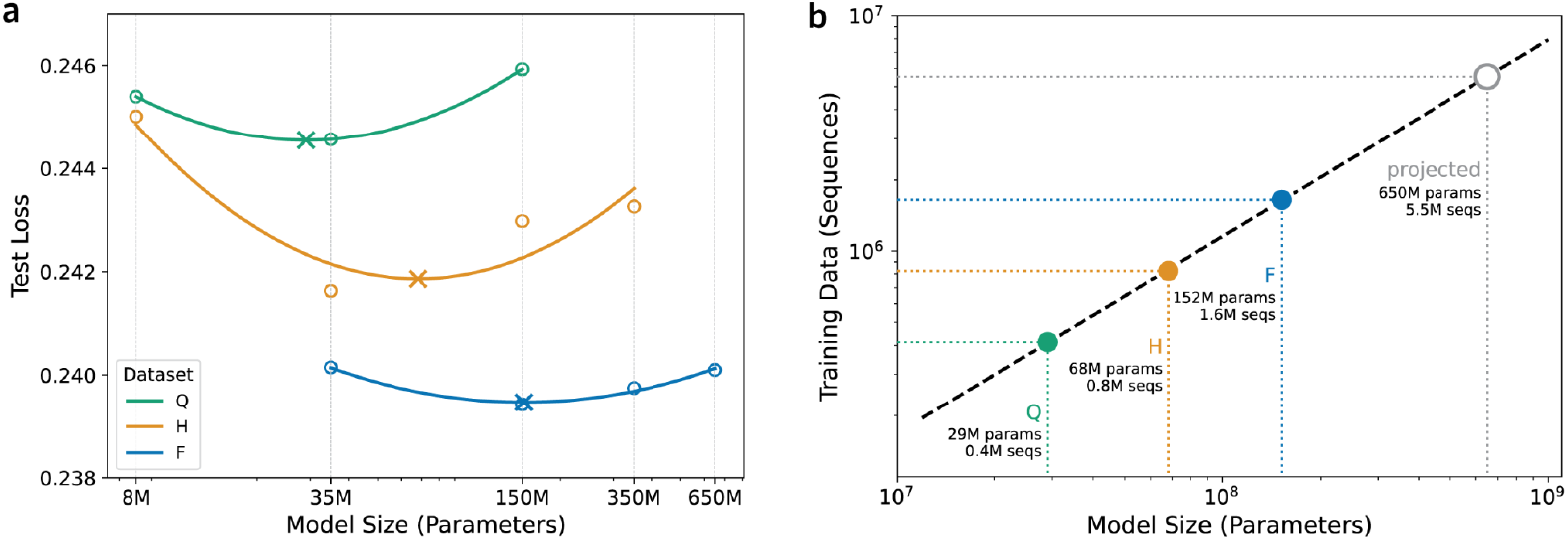
Establishing FixedData profiles: (a) For each dataset size, model sizes were log-transformed to linearize the exponentially scaling relationship with evaluation loss. A quadratic regression was then fit to the transformed data to create FixedData profiles (Dataset Q, R^2^ = 1.00; Dataset H, R^2^ = 0.84; Dataset F, R^2^ = 0.98), with the x-axis displayed on a logarithmic scale. Circles (O) represent the mean loss averaged over 10 donor datasets for each model–data size combination, while crosses (X) mark the minima of the fitted curves, corresponding to the model size achieving the lowest loss. (b) Using the inferred optimal model sizes from (a), we estimate the optimal data size for a 650M-parameter AbLM by fitting a power-law regression model (R^2^ = 0.9997).

The 650M parameter variant of the ESM-2 protein LM is highly performant and widely used (6); we estimate that data-optimal training of a similarly sized AbLM will require approximately 5.5M paired antibody sequences (***Figure 2b***), which is roughly double the amount of paired antibody sequences currently present in the Observed Antibody Space (OAS) database (29).

### Optimally scaled AbLMs improve residue identity prediction primarily in the highly variable CDRH3 regions

To explore if models deemed optimal by FixedData profiles (which are based on MLM loss) also perform the best on tasks that better represent real-world use cases, we evaluated the models on a series of benchmarks. First, we sampled 1,000 mutated and unmutated sequences from each of the 10 evaluation datasets and assessed all models pretrained using Dataset-F for their ability to predict masked residues. Heavy chain sequences were iteratively masked, and for each antibody region we calculated the median cross-entropy loss across all masked positions in that region for mutated and non-mutated sequences (***Figure 3, Table S4***). All models show comparable performance in framework regions (FWRs), presumably due to the inherent germline bias of most AbLMs (21), but differed more on non-templated heavy chain CDR3s. All model sizes perform similarly on unmutated CDR3 (***Figure 3a***), but the 150M-F and 350M-F models outperform other models on the mutated CDR3 (***Figure 3b***), in agreement with the FixedData-based prediction that the optimal parameter count falls between these two model sizes. 650M-F performed worse than both 150M-F and 350M-F, likely because 650M-F overfit more rapidly than the other models, underscoring the importance of training optimally sized models.

**Figure 3.**
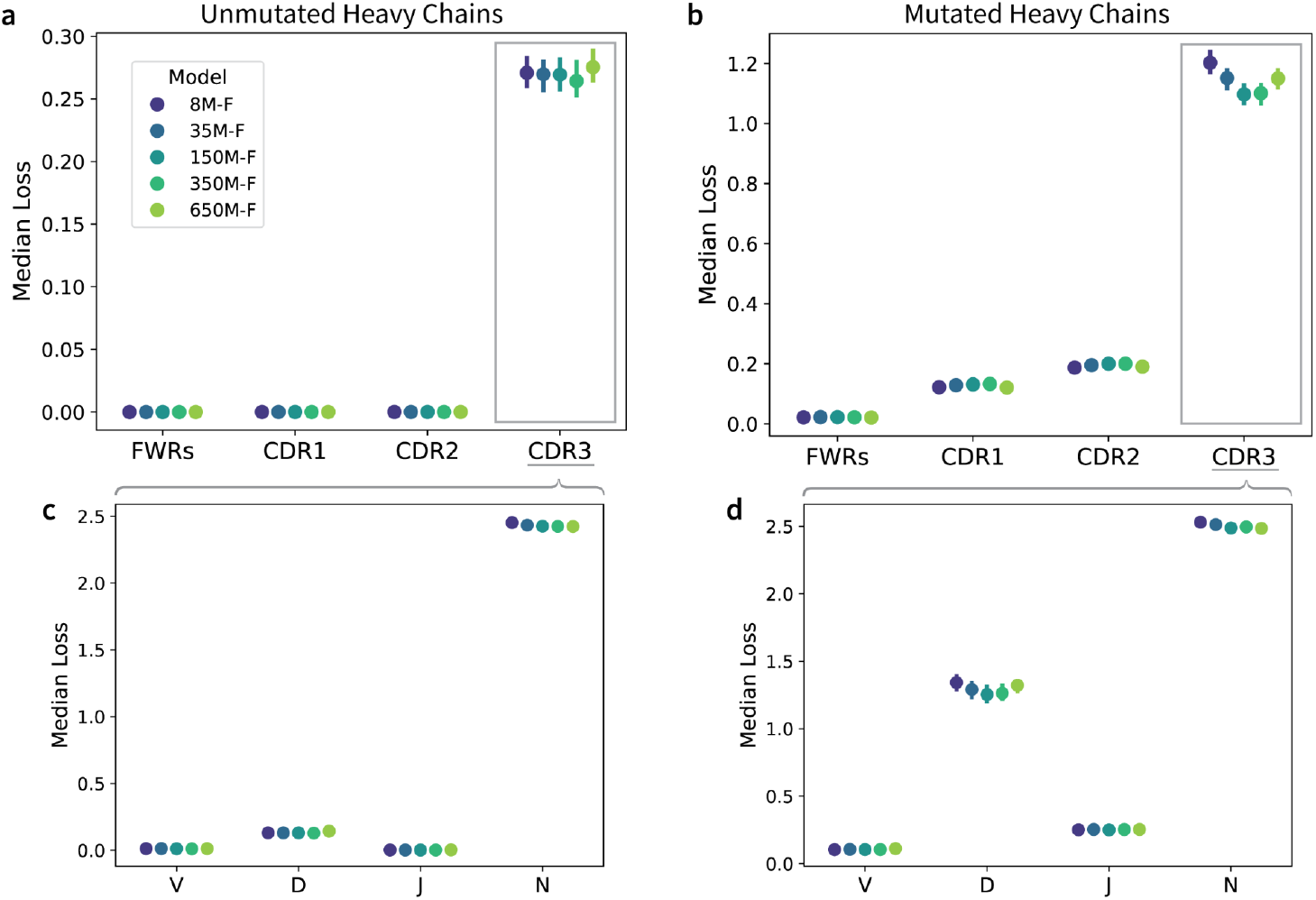
Regional per-residue prediction performance. Median per-residue cross-entropy loss for each AbLM trained on Dataset-F across four antibody regions in the heavy chain (FWRs, CDRH1–3), evaluated separately on 10,000 unmutated (a) and mutated (b) antibody sequences from 10 unique donors. Predictions in the CDRH3 are further broken down into subregions, including V, D, and J gene-derived segments, and non-templated N-addition residues for unmutated (c) and mutated (d) sequences. Error bars represent the 95% confidence interval (CI) of the median.

To further examine model performance in the CDRH3, we partitioned CDRH3 residues based on their derivation from V, D, and J gene segments or from the non-templated (N-addition) regions generated during V(D)J recombination (***Figure 3c-d***). We observe that the median loss is substantially higher for N-addition residues of both unmutated and mutated sequences, as expected due to the increased sequence variability introduced during junctional diversification. As model size increases from 8M to 350M parameters, we observe a clear reduction in median loss on D gene residues in mutated sequences. Paired t-tests across model sizes (***Table S5***) show significant differences on D gene predictions for 8M vs 35M (p=0.001), 8M vs 150M (p=0.017), 8M vs 350M (p=0.017), and 35M vs 650M (p=0.030). Because the D gene contributes the largest share of templated residues in CDRH3, these trends likely contribute significantly to the aggregate model performance across the CDRH3 region. This suggests that larger, data-optimal AbLMs start to learn the D gene region in mutated sequences but not the stochastic patterns in non-templated regions.

### Joint scaling of data and model size improves specificity classification performance

Next, we assessed model performance on binary (coronavirus (CoV)-specific or healthy donor) (***Figure 4a, c***) and multi-class (CoV-specific, influenza-specific, or healthy donor) (***Figure 4b, d***) antibody specificity classification using 5-fold cross-validation (CV). Plotting the binary classification accuracy of each model reveals an interesting scaling phenomenon (***Figure 4a***). All models exhibit better performance as the training data scale increases, except the 8M parameter model. The 8M parameter models perform similarly regardless of training data scale, consistent with our previous observation that 8M parameter models are likely too small to benefit from additional pretraining data. We see the largest performance separation at 350M parameters, where 350M-F significantly outperforms 350M-H. However, we observe no further improvements in classification performance when the models are scaled up to 650M parameters. Similar trends are observed across the other classification metrics (***Figure 4c, Table S6***).

**Figure 4.**
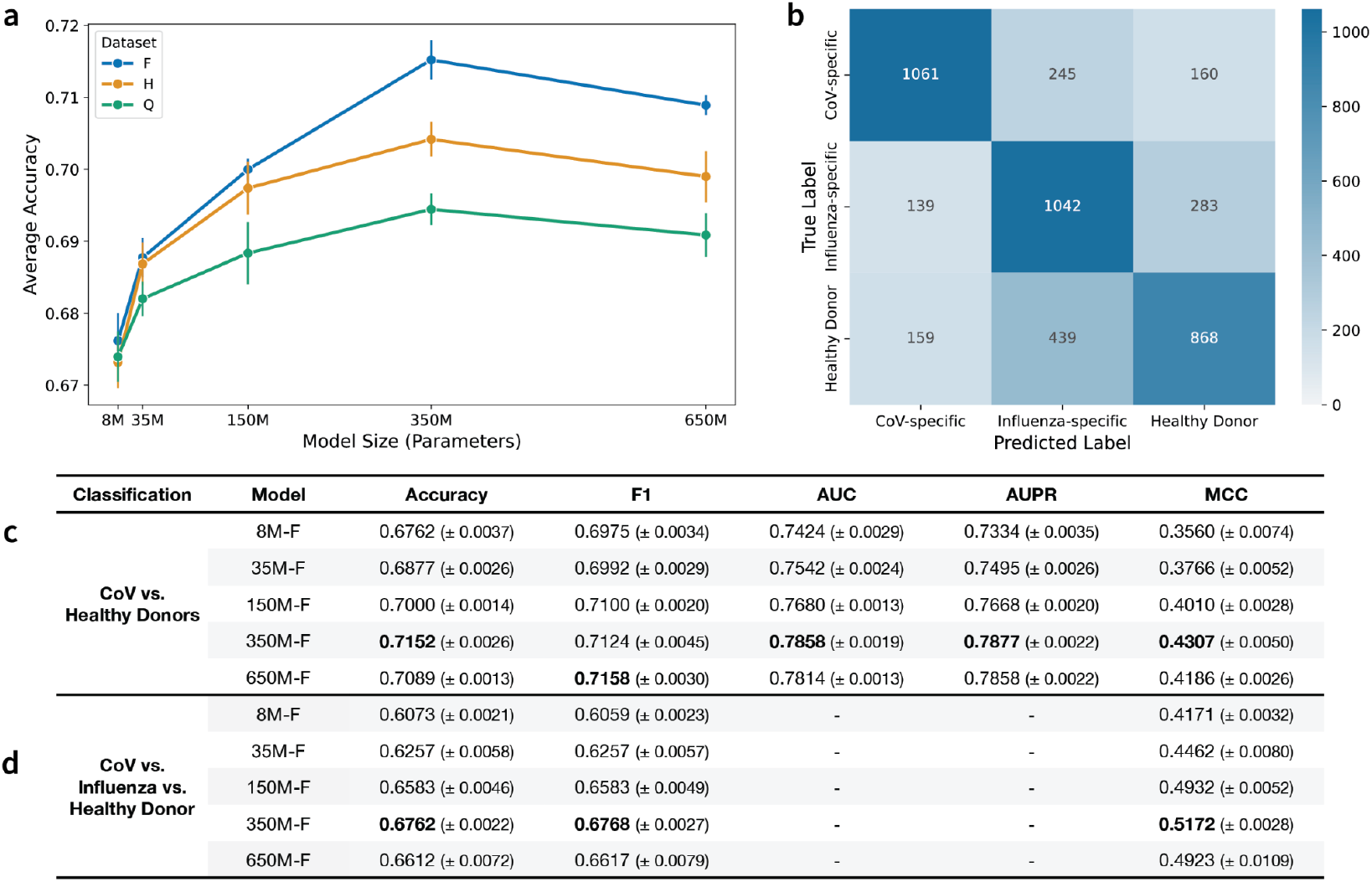
Antibody specificity classification performance across the dataset and model sizes. (a) Average binary classification accuracy as a function of model size and dataset size. Error bars denote the standard error across 5-fold CV replicates. (b) Confusion matrix for the three-way classifier model trained on the 350M-F base. (c) Binary classification results distinguishing CoV-specific antibodies and Healthy Donor antibodies for Dataset-F models. (d) Three-way classification results distinguishing CoV-specific, Influenza-specific, and Healthy Donor antibodies for Dataset-F models. For each classification task, the best overall model for each metric is indicated in **bold**.

In the multi-class task, the 350M-F parameter model performed the best across the classification metrics (***Figure 4d***). To further analyze the predictions of the 350-F model, we plot a confusion matrix comparing the true and predicted classes across all 5 folds (***Figure 4b***). The confusion matrix reveals strong diagonal dominance, with the majority of sequences correctly classified into their respective categories. However, the model frequently misclassifies healthy donor sequences as influenza-specific, likely due to the construction of the classification dataset: the ‘healthy donor’ repertoires may contain influenza-specific antibodies, making it difficult for the model to separate the two classes.

Overall, the 350M-F model consistently outperforms the other models across model sizes and data scales on both classification tasks (***Table S6***). This suggests that the FixedData profiles may slightly underestimate the performance-optimal model size, particularly on tasks that differ from the MLM objective used to compute test loss. In addition, the reduction of performance observed in the 650M models suggests that in data-limited regimes like natively paired AbLMs, increasing the available training data is a prerequisite for scaling model size beyond existing thresholds.

### Data-optimal AbLMs improve recognition of natively paired antibody chains

Based on mounting evidence that antibody heavy and light chain pairing is not entirely random (39), we and others have previously fine-tuned AbLMs to distinguish between natively paired and randomly paired antibody heavy and light chains (23,40). To assess the effects of scaling on this task, we fine-tuned each of our models to perform a binary classification of whether a paired sequence is a native or shuffled pair with 5-fold CV (***Figure 5, Table S7***). Even the smallest models achieve a level of classification accuracy that surpasses random guessing, but improvement over this baseline accuracy is not observed until the model size reaches 350M parameters (***Figure 5a***). In line with previous results, we additionally observe that the 650M parameter models consistently perform worse than the 350M models. 350M parameters also represents the model size for which classification performance across the three training data scales becomes distinguishable. Further prediction outcome analysis of the 350M parameter models revealed that performance gains are driven by improved classification of native pairs, while shuffled pair classification remained largely unchanged across increasing data scales (***Figure 5b***). This suggests that as models become more data-optimal, features that signal native chain pairing become more apparent in the resulting sequence embeddings.

**Figure 5.**
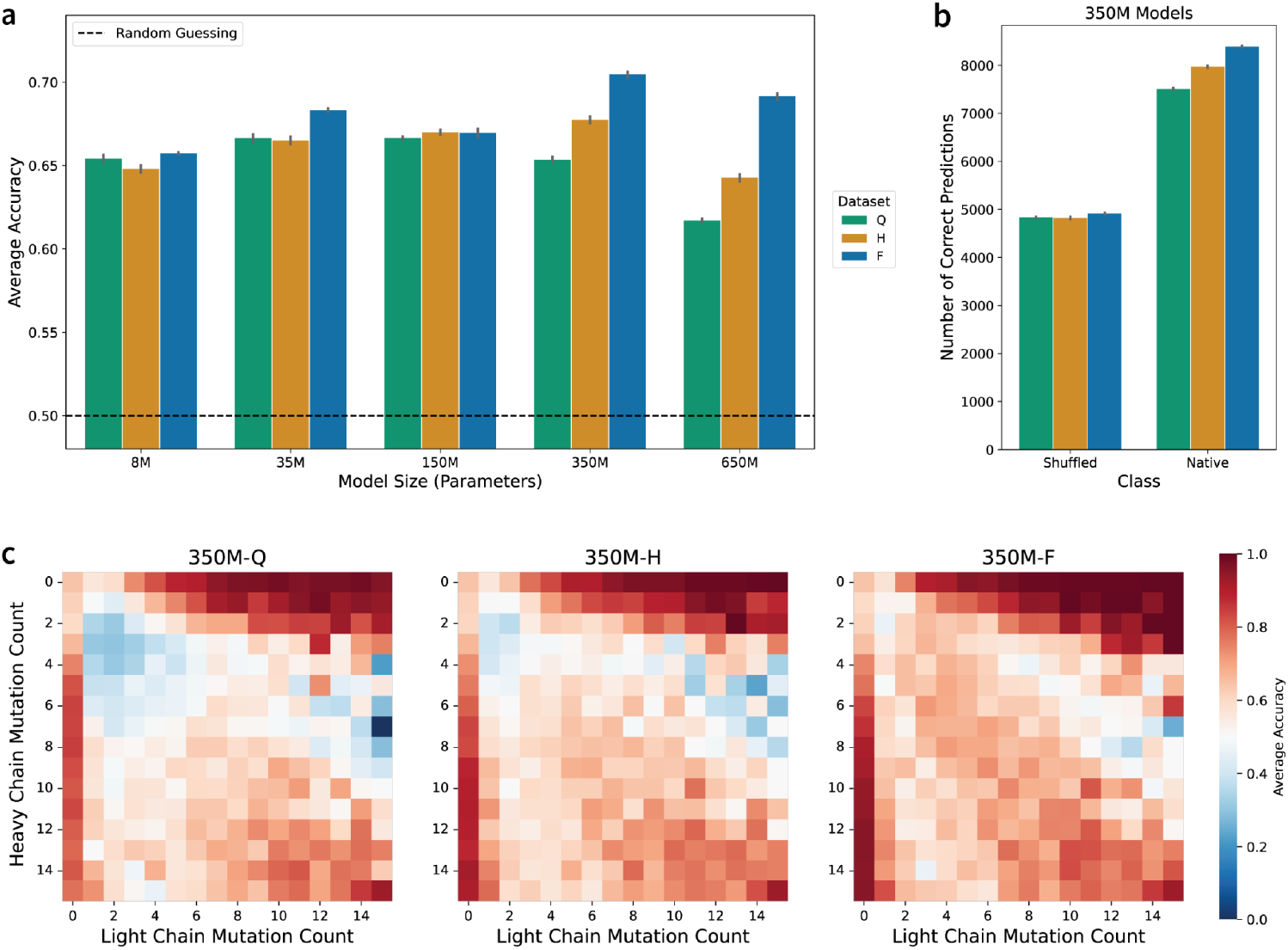
Comparison of model performance on the native vs. shuffled chain pairing classification task. (a) Average classification accuracy for all model sizes and data scales. (b) Number of correct predictions of the shuffled and native classes for the 350M model classifiers. Error bars indicate the standard error of the mean across 5-fold CV replicates. (c) Average classification accuracy by the 350M model classifiers for different combinations of chain-specific mutation counts. More accurate predictions are darker red.

Previous studies have demonstrated that much of the accuracy achieved by current pair classification models results from learning that natively paired chains tend to have similar levels of somatic hypermutation (23). To identify whether a similar heuristic explains our observed results, we analyzed the performance of the 350M models across different combinations of chain-specific mutation counts (***Figure 5c***). As data scale increases, improvements tend to be focused on sequence pairs with low and similar mutation counts (1-7 mutations) in both the heavy and light chains: average prediction accuracy in these pairs was observed to increase from 45.9% in Dataset-Q, to 54.2% in Dataset-H, to 63.2% in Dataset-F. Since performance gains are primarily observed in the correct classification of native pairs with similar mutation counts, our models are likely using the chain-specific mutation count heuristic to correctly identify native pairs more frequently. This heuristic must be learned implicitly, as mutation count information is not explicitly provided during training (23). We similarly observe improved performance within other model sizes (***Figure S3***), suggesting that increased data scale promotes improved learning of affinity maturation-driven mutation.

## DISCUSSION

Existing scaling laws for NLP and protein LMs focus on compute-optimal scaling (31), given that these models are primarily compute-constrained. Here, we explore the optimal scaling of natively paired AbLMs, where the primary limitation is the amount of available training data. We trained 15 AbLMs to explore the relationship between model size and training data scale.

FixedData profiles revealed that the optimal training data volume scales as a power law of model size. The 650M parameter variant of the ESM-2 protein LM is widely used because it is both highly performant and compatible with the GPUs commonly found in academic research environments. By extrapolation, we show that optimal training of a 650M-parameter AbLM will require ∼5.5M sequences, which is approximately twice the number of paired antibody sequences currently available in the Observed Antibody Space repository (28,29). Notably, these figures correspond to the optimal number of sequences after clustering at 90% identity; therefore, the total number of paired sequences required will likely be higher.

One major area of concern for AbLMs is their poor performance on the highly variable, but functionally critical, CDR3 loops. To further evaluate this poor performance, we assessed the model’s ability to predict the V, D, J, and N addition regions of the CDRH3. We observe very little improvement in model performance on N addition regions, which is expected given that N additions are stochastic and drive much of the antibody repertoire’s diversity (41). However, the optimally scaled 150M-F and 350M-F models show improved performance at predicting mutated D gene regions.

Additionally, each of the pretrained AbLMs was evaluated on a suite of downstream tasks designed to more accurately mimic real-world use cases. The best performing models on these tasks were typically larger than the size predicted by FixedData profiles (∼152M parameters for Dataset-F). Specifically, the 350M parameter models consistently outperformed the 150M parameter models on downstream classification tasks. One potential explanation for this observation is the size of the output projection layer used for the classification tasks. The 350M model has a larger hidden dimension than the 150M model, which results in more trainable parameters in its classification head that are not accounted for by FixedData. However, this factor alone does not fully explain performance differences, as the 650M model has the largest output layer but is not the top performer. This suggests that when the optimal model size falls between two feasible model sizes, selecting the larger of the two sizes may be a viable strategy, but scaling too far beyond optimal will lead to deteriorated performance.

The results of our pairing classification task indicate that a subset of randomly paired chains with mismatched mutation frequencies is easily distinguishable by even the smallest models, but the remaining examples require more sophistication to classify accurately. This suggests that pairing classification may become an increasingly usable function of AbLMs as they are scaled with sufficient data. The potential for emergent abilities (42), which appear in NLP models as model size increases, may be observed in AbLMs as model and data sizes are optimally scaled.

While not explored here, previous studies have implemented methods for improving data efficiency in AbLMs, such as focal loss or preferential masking (21,22). When applied effectively, incorporating these methods into pretraining would alter the scaling law such that fewer paired sequences are required to achieve a data-optimal 650M parameter model. This is an interesting potential direction for future studies, given the limitations that prevent rapid generation of large paired sequence datasets. However, data efficiency methods do not overcome the need for more paired sequencing data, and it remains important to consider the balance between model size and data scale.

Our work highlights the current data bottlenecks for specific downstream tasks and provides practical guidelines for estimating the amount of data required to optimally train AbLMs of varying sizes. By establishing clear scaling laws, we offer a systematic approach to balancing data collection efforts against model complexity. Ultimately, our results emphasize that future improvements in antibody language modeling will increasingly depend on dedicated efforts to expand and diversify paired antibody datasets to expand the downstream capabilities of future AbLMs.

## METHODS

### Datasets

The pretraining data was downloaded from the OAS (29) on September 12th, 2024, and supplemented with sequences from Jaffe et al. (39) and an internally generated dataset of ∼400k sequences from healthy donor B cells. These sequences were derived from circulating B cells of healthy adult donors without any selection or enrichment for binding to a specific antigen. Raw sequences were annotated using abstar (43), filtered as described in AntiRef (44), and clustered at 90% identity using MMseqs (45), resulting in 1,717,423 sequence pairs.

For model pretraining, we partitioned the full dataset such that 96% was allocated for training (1,648,726 pairs), while 2% was held out for evaluation (34,349 pairs) and an additional 2% for testing. To investigate the influence of dataset size on model performance, we further derived two training subsets from the primary training set. First, we randomly selected 50% of the training pairs (824,363 pairs) to create a half-size dataset, from which a further random selection of 50% (412,182 pairs) was made to form the quarter-size dataset. Paired sequences were concatenated with two <cls> tokens as the separator and tokenized using the ESM-2 tokenizer (6), with a vocabulary of 33 tokens.

For model evaluation, we used an internally generated collection of paired antibody sequence datasets from 10 distinct donors not present in the training set. This reduces the impact of donor-specific effects from pretraining, ensuring the generalizability of our findings. Sequences were clustered at 90% identity, resulting in 94,483 paired antibody sequences.

### Model pretraining

We trained fifteen ESM-2 architecture (6) language models of varying sizes (approximately 8, 35, 150, 350, and 650 million parameters) on the three paired antibody datasets described above. The model parameters are provided in more detail in ***Table S1***.

Models were trained with the HuggingFace Transformers library (46), using a masked language modeling objective. For each training sequence, 15% of the sequence was randomly selected for prediction, and of these, 80% were masked, 10% were replaced with a random token, and 10% were left unchanged. Models were trained for 500,000 steps, with a linear warm-up of 30,000 steps, and a peak learning rate of 1×10−^4^. The total batch size was 128 per update, trained on 4 GPUs. We ensured reproducibility by setting a random seed of 42. Training progress and metrics were logged using Weights & Biases (47).

### FixedData Evaluation

To calculate the optimal model size for each dataset size, we used our 10 distinct donor test datasets and adopted the approach used by (31) to construct “FixedData profiles”. We retain only the points around the minimum for each curve for quadratic regression analysis; including points far from this minimum would dilute the estimate of the optimal configuration and introduce unnecessary variance.

To investigate the relationship between model performance (test loss), model size (parameter count), and pretraining dataset scale, we modeled average test loss as a quadratic function of the log-transformed parameter counts using the following equation:

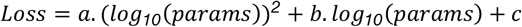

The minimum of each FixedData curve was used to determine the optimal model sizes for each dataset scale. We then projected the optimal amount of training data for an ESM-2-sized AbLM (650M parameters) by fitting a power-law regression to these FixedData minima (***Figure 2B***). We obtained the following optimal scaling relationship with strong fit (*R*^*2*^ = 0.*9996*) :

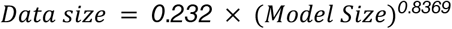

This scaling law indicates sub-linear scaling, suggesting diminishing marginal data requirements as models grow and providing practical guidance for efficient model scaling strategies.

*Classification tasks*. For the specificity classification tasks, the datasets consisted of 27,442 paired sequences (13,721 for each class) for the binary classification task (healthy donor vs CoV) and 4,398 paired sequences (1,466 for each class) for the three-way classification task (healthy donor vs CoV vs Flu). CoV-specific sequences were sourced from the CoV-AbDab (48), Flu-specific antibodies were obtained from Wang et al. (49), and healthy donor antibodies were obtained from the Ng et al. control dataset (23). For the binary classification, models were fine-tuned (with the base model weights frozen) for 3 epochs with a batch size of 128. For the multi-class classification, models were fine-tuned for 5 epochs with a batch size of 32.

For the native pairing classification task, we shuffled our 10 distinct donor test datasets as described in (23). Shuffled pairs were generated by randomly sampling 50% of the sequences from each donor and shuffling their heavy and light chains. In total, the dataset comprised 94,414 antibody sequence pairs (47,207 for each class). The models were fine-tuned for 50 epochs with a batch size of 256.

All classification tasks were performed using 5-fold CV with stratification and different random seeds during training. All models were trained using a linear learning rate scheduler with a 10% warmup ratio and a peak learning rate of 5 × 10−^5^ To evaluate the classifier performance, we computed several metrics: accuracy, F1 score, area under the receiver operating characteristic curve (AUC), area under the precision-recall curve (AUPR), and Matthews correlation coefficient (MCC).

### Code and Data Availability

All code used for data processing, model training, and evaluation is available at this GitHub repository: https://github.com/brineylab/AbLMs-scaling-laws/.

Pretrained AbLMs and associated model checkpoints are archived and openly accessible through Zenodo at https://zenodo.org/records/16938681. All models are also hosted on the HuggingFace Model Hub at https://huggingface.co/collections/brineylab/ablms-scaling-laws-6824e4beaabf4b16107cac4f, where users can load and fine-tune the models using standard HuggingFace tools.

## AUTHOR CONTRIBUTIONS

B.B. and M.S.N conceptualized the study. Model training and evaluation were carried out by M.S.N., S.B., K.N., and B.B. Data generation was performed by J.H., J.M., N.I., D.M, and T.N. The manuscript was prepared, revised and reviewed by all authors.

## ACKNOWLEDGMENTS

The authors would like to thank Monica L. Fernández-Quintero and Johannes Loeffler for assistance with manuscript revisions; Terrence Messmer Jr. and Charles Bowman for facilitating access to compute resources for model training; and Simone Spandau for support in generating sequencing data.

## SUPPLEMENTARY INFORMATION

**Table S1.** Model architecture details and checkpoint selections. Each row corresponds to a distinct configuration of a pretrained model, varying by model size (in millions of parameters), training data size (Full, Half, or Quarter), and chosen checkpoint (in training steps).

| Model size (M) | Dataset | Checkpoint step | Transformer layers | Attention heads | Hidden size | Intermediate size |
| --- | --- | --- | --- | --- | --- | --- |
| 8 | F | 500,000 | 6 | 20 | 320 | 1280 |
|  | H | 435,000 |  |  |  |  |
|  | Q | 425,000 |  |  |  |  |
| 35 | F | 500,000 | 12 | 20 | 480 | 1920 |
|  | H | 430,000 |  |  |  |  |
|  | Q | 240,000 |  |  |  |  |
| 150 | F | 500,000 | 30 | 20 | 640 | 2560 |
|  | H | 330,000 |  |  |  |  |
|  | Q | 165,000 |  |  |  |  |
| 350 | F | 500,000 | 32 | 20 | 960 | 3840 |
|  | H | 300,000 |  |  |  |  |
|  | Q | 155,000 |  |  |  |  |
| 650 | F | 395,000 | 33 | 20 | 1280 | 5120 |
|  | H | 330,000 |  |  |  |  |
|  | Q | 130,000 |  |  |  |  |

**Figure S1.**
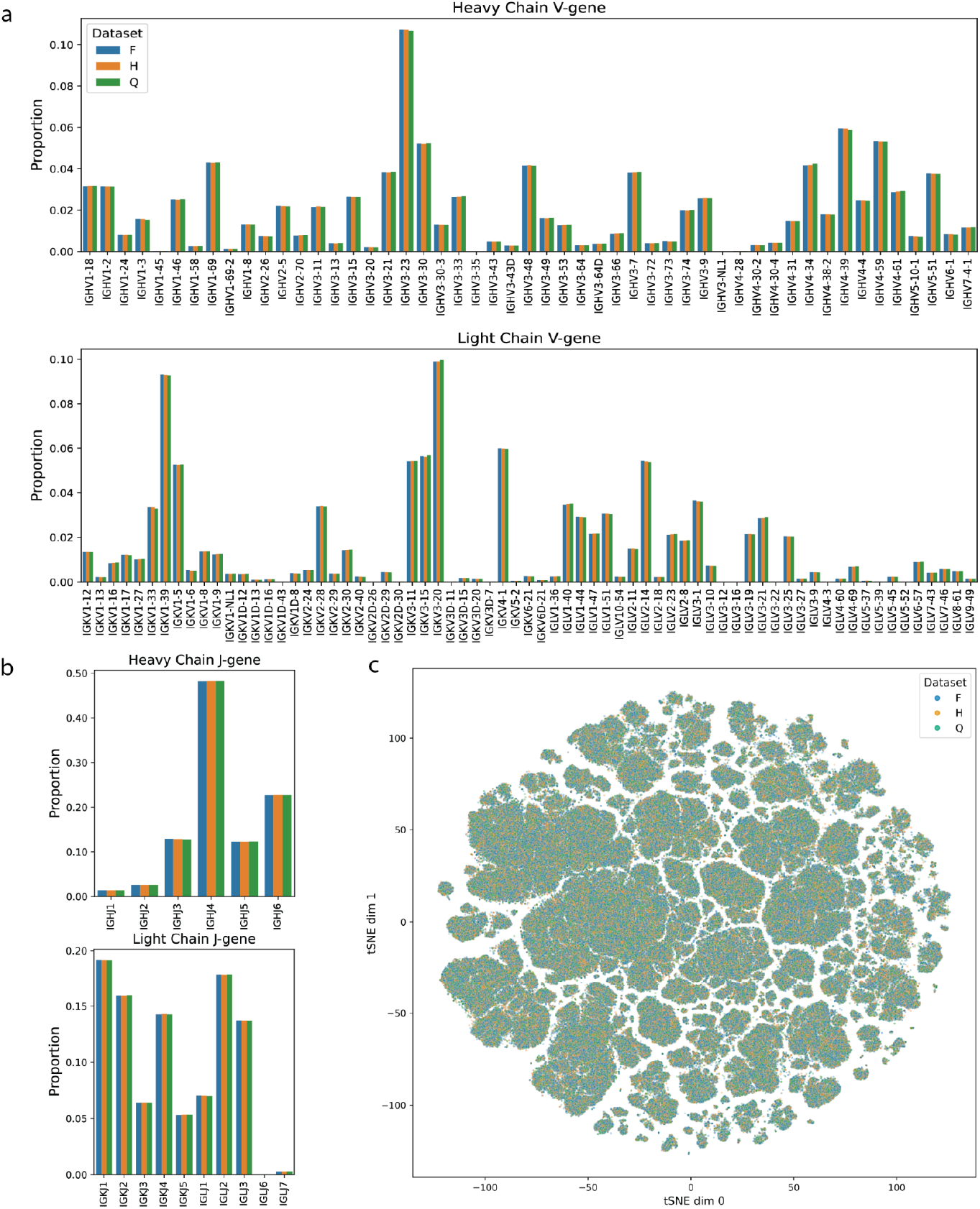
Distribution of V- and J-gene usage and sequence diversity across datasets. (a) V-gene usage across training-set splits. Grouped bar plots show the proportion of sequences in each dataset with a particular heavy-chain V (top) or light-chain V (bottom) gene. (b) J-gene usage across training-set splits. Grouped bar plots show the proportion of sequences in each dataset with a particular heavy-chain J (top) or light-chain J (bottom) gene. (c) Sequence diversity visualized using a t-SNE projection of ESM-2-650M embeddings, with each point representing a single paired sequence. All training sets exhibit comparable diversity.

**Table S2.** χ^2^ tests of gene-usage distributions across training-set splits. Pairwise comparisons between datasets (Q vs H, H vs F, Q vs F) of heavy and light chain V, J, and V/J gene-usage distribution. Comparisons are reported with the number of categories tested, the χ^2^ statistic, and the corresponding p-value.

| Feature | Training Splits | # of Categories | $\chi^2$ Statistic | $\chi^2$ p-value |
| --- | --- | --- | --- | --- |
| Heavy Chain V-gene | Quarter vs Half | 51 | 20.079451 | 0.999950 |
|  | Half vs Full | 51 | 17.889161 | 0.999992 |
|  | Quarter vs Full | 51 | 37.792155 | 0.897789 |
| Heavy Chain J-gene | Quarter vs Half | 6 | 1.066540 | 0.957022 |
|  | Half vs Full | 6 | 2.090143 | 0.836537 |
|  | Quarter vs Full | 6 | 3.341937 | 0.647428 |
| Heavy Chain VJ-gene usage | Quarter vs Half | 299 | 107.238132 | 1.000000 |
|  | Half vs Full | 301 | 101.794289 | 1.000000 |
|  | Quarter vs Full | 301 | 198.955262 | 0.999999 |
| Light Chain V-gene | Quarter vs Half | 69 | 22.444984 | 1.000000 |
|  | Half vs Full | 69 | 25.491875 | 0.999999 |
|  | Quarter vs Full | 69 | 44.962355 | 0.986015 |
| Light Chain J-gene | Quarter vs Half | 10 | 2.113995 | 0.989534 |
|  | Half vs Full | 10 | 1.188426 | 0.998866 |
|  | Quarter vs Full | 10 | 4.035732 | 0.909045 |
| Light Chain VJ-gene usage | Quarter vs Half | 302 | 94.901586 | 1.000000 |
|  | Half vs Full | 304 | 100.277863 | 1.000000 |
|  | Quarter vs Full | 304 | 179.888776 | 1.000000 |

**Figure S2.**
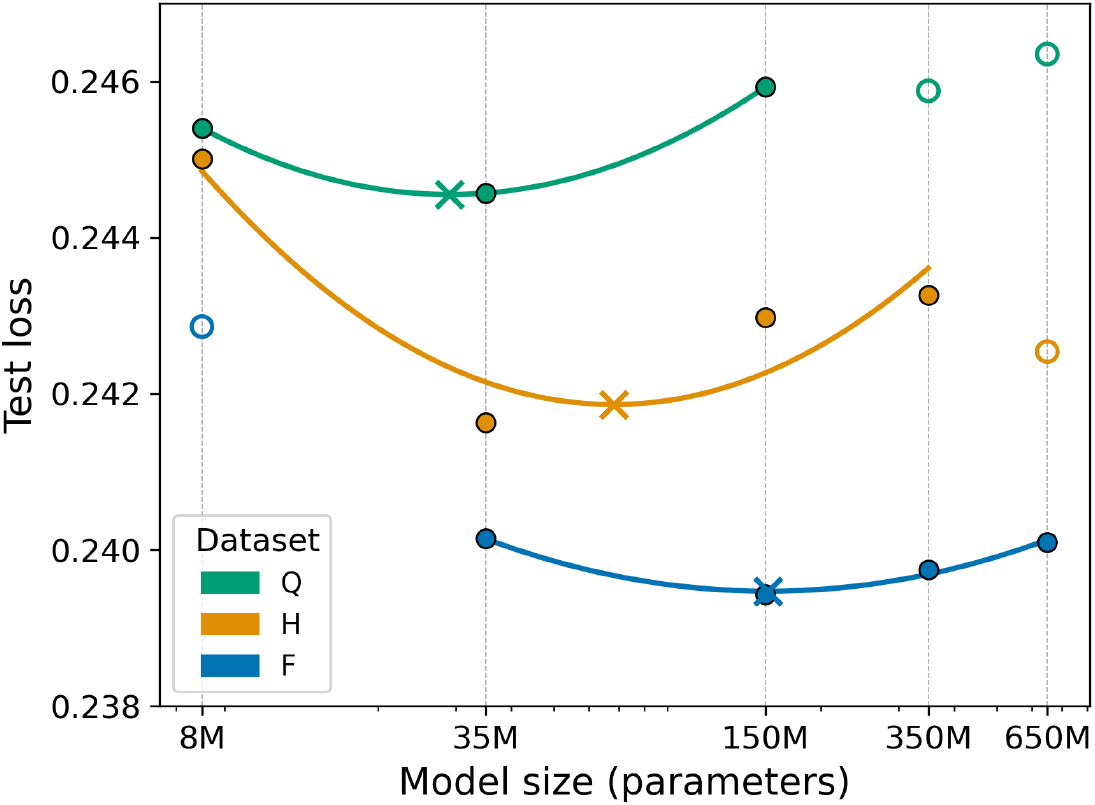
Cross-entropy loss curves with all evaluated points for each dataset size. Model sizes were log-transformed and evaluated to generate FixedData profiles. Circles (O) represent the mean loss averaged over 10 donor datasets for each model–data size combination, while crosses (X) mark the fitted curve minimum corresponding to the lowest loss. In this figure, all evaluated mean cross entropy losses are displayed, with outlier points shown as unfilled circles.

**Table S3.** Average cross-entropy loss during evaluating models across different scales that were evaluated using a masked language modeling (MLM) objective on data from 10 distinct donors.

| Dataset | Model size (M) | Average loss |
| --- | --- | --- |
| Q | 8 | 0.2454 |
|  | 35 | 0.2445 |
|  | 150 | 0.2459 |
|  | 350 | 0.2458 |
|  | 650 | 0.2463 |
| H | 8 | 0.2450 |
|  | 35 | 0.2416 |
|  | 150 | 0.2429 |
|  | 350 | 0.2432 |
|  | 650 | 0.2425 |
| F | 8 | 0.2428 |
|  | 35 | 0.2401 |
|  | 150 | 0.2394 |
|  | 350 | 0.2397 |
|  | 650 | 0.2401 |

**Table S4.** CDRH3 prediction metrics for all full data models. Median cross entropy loss and median perplexity of models trained on the full dataset for per-residue prediction in the CDRH3. Metrics are shown across varying model sizes (in millions of parameters) and grouped by mutated and germline (unmutated) sequence categories.

| Region | Condition | Model size (M) | Loss | Perplexity |
| --- | --- | --- | --- | --- |
| CDRH3 | Mutated | 8 | 1.2026 | 3.4237 |
|  |  | 35 | 1.1513 | 3.2293 |
|  |  | 150 | 1.0970 | 3.0614 |
|  |  | 350 | 1.1009 | 3.0585 |
|  |  | 650 | 1.1502 | 3.2378 |
|  | Unmutated | 8 | 0.2709 | 1.3149 |
|  |  | 35 | 0.2797 | 1.3152 |
|  |  | 150 | 0.2694 | 1.3136 |
|  |  | 350 | 0.2643 | 1.3092 |
|  |  | 650 | 0.2754 | 1.3239 |

**Table S5.**
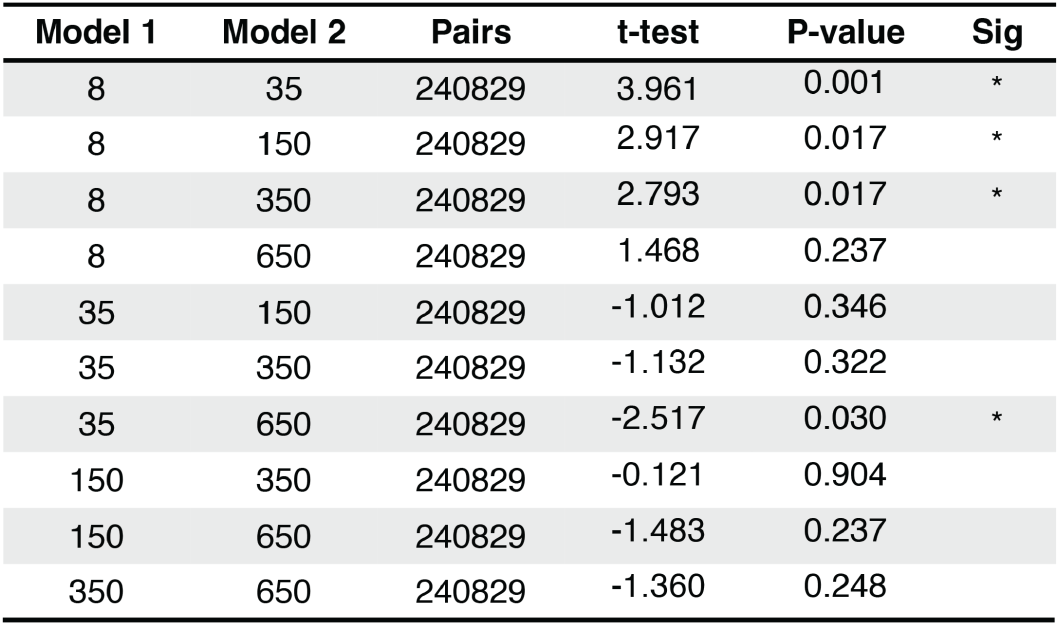
Paired t-test across model sizes for D-segment prediction on the CDRH3 region. Each comparison reports the t-statistic, Benjamini–Hochberg corrected p-value, and significance level for paired evaluations across models. Asterisks indicate statistically significant differences between models.

| Model 1 | Model 2 | Pairs | t-test | P-value | Sig |
| --- | --- | --- | --- | --- | --- |
| 8 | 35 | 240829 | 3.961 | 0.001 | * |
| 8 | 150 | 240829 | 2.917 | 0.017 | * |
| 8 | 350 | 240829 | 2.793 | 0.017 | * |
| 8 | 650 | 240829 | 1.468 | 0.237 |  |
| 35 | 150 | 240829 | -1.012 | 0.346 |  |
| 35 | 350 | 240829 | -1.132 | 0.322 |  |
| 35 | 650 | 240829 | -2.517 | 0.030 | * |
| 150 | 350 | 240829 | -0.121 | 0.904 |  |
| 150 | 650 | 240829 | -1.483 | 0.237 |  |
| 350 | 650 | 240829 | -1.360 | 0.248 |  |

**Table S6:** Performance of antibody specificity classification models across AbLMs trained on Dataset-H and Dataset-Q. Binary classification results for distinguishing CoV-specific antibodies from healthy donor sequences. Three-way classification results differentiating Influenza-specific, CoV-specific, and healthy donor antibodies. For each dataset size, the best model is indicated in **bold** per metric.

| Classification | Dataset | Model size (M) | Accuracy | AUC | AUPR | MCC | F1 Score |
| --- | --- | --- | --- | --- | --- | --- | --- |
| CoV vs. Healthy Donors | Quarter | 8 | 0.6740 (± 0.0034) | 0.7362 (± 0.0025) | 0.7253 (± 0.0024) | 0.3504 (± 0.0067) | 0.6923 (± 0.0028) |
| | | 35 | 0.6820 (± 0.0023) | 0.7445 (± 0.0024) | 0.7354 (± 0.0018) | 0.3643 (± 0.0045) | 0.6866 (± 0.0026) |
| | | 150 | 0.6884 (± 0.0042) | 0.7541 (± 0.0037) | 0.7520 (± 0.0037) | 0.3773 (± 0.0083) | 0.6954 (± 0.0046) |
| | | 350 | **0.6944** (± 0.0021) | **0.7673** (± 0.0015) | **0.7699** (± 0.0016) | **0.3900** (± 0.0041) | **0.7048** (± 0.0025) |
| | | 650 | 0.6909 (± 0.0029) | 0.7608 (± 0.0020) | 0.7656 (± 0.0016) | 0.3827 (± 0.0059) | 0.6981 (± 0.0053) |
| | Half | 8 | 0.6732 (± 0.0035) | 0.7411 (± 0.0031) | 0.7350 (± 0.0033) | 0.3520 (± 0.0071) | 0.6998 (± 0.0032) |
| | | 35 | 0.6869 (± 0.0028) | 0.7544 (± 0.0025) | 0.7487 (± 0.0038) | 0.3756 (± 0.0057) | 0.7014 (± 0.0033) |
| | | 150 | 0.6974 (± 0.0036) | 0.7667 (± 0.0023) | 0.7667 (± 0.0018) | 0.3955 (± 0.0072) | 0.7057 (± 0.0043) |
| | | 350 | **0.7042** (± 0.0023) | **0.7765** (± 0.0017) | **0.7818** (± 0.0027) | **0.4088** (± 0.0045) | **0.7093** (± 0.0033) |
| | | 650 | 0.6990 (± 0.0034) | 0.7697 (± 0.0023) | 0.7716 (± 0.0031) | 0.3988 (± 0.0068) | 0.7067 (± 0.0043) |
| Influenza vs. CoV vs. Healthy Donor | Quarter | 8 | 0.6062 (± 0.0051) | - | - | 0.4169 (± 0.0067) | 0.6058 (± 0.0049) |
| | | 35 | 0.5769 (± 0.0109) | - | - | 0.3699 (± 0.0159) | 0.5751 (± 0.0119) |
| | | 150 | 0.6444 (± 0.0078) | - | - | 0.4694 (± 0.0118) | 0.6448 (± 0.0074) |
| | | 350 | **0.6467** (± 0.0057) | - | - | **0.4731** (± 0.0076) | **0.6462** (± 0.0059) |
| | | 650 | 0.6353 (± 0.0041) | - | - | 0.4558 (± 0.0062) | 0.6364 (± 0.0036) |
| | Half | 8 | 0.5928 (± 0.0055) | - | - | 0.3986 (± 0.0056) | 0.5894 (± 0.0060) |
| | | 35 | 0.6253 (± 0.0083) | - | - | 0.4458 (± 0.0116) | 0.6230 (± 0.0093) |
| | | 150 | 0.6476 (± 0.0084) | - | - | 0.4756 (± 0.0117) | 0.6481 (± 0.0090) |
| | | 350 | **0.6610** (± 0.0043) | - | - | **0.4949** (± 0.0067) | **0.6619** (± 0.0038) |
| | | 650 | 0.6576 (± 0.0026) | - | - | 0.4888 (± 0.0036) | 0.6597 (± 0.0026) |

**Table S7.** Detailed classification results for pair classification. Performance metrics are presented for models ranging from 8M to 650M parameters, evaluated across Dataset-F, Dataset-H, and Dataset-Q. The best-performing model for each metric across all datasets is shown in **bold**, while the top-performing model for each metric within individual datasets is underlined.

| Classification | Dataset | Model Size (M) | Accuracy | AUC | AUPR | MCC | F1 Score |
| --- | --- | --- | --- | --- | --- | --- | --- |
| Native vs. Shuffled Pairing | Quarter | 8 | 0.6540 (± 0.0024) | 0.6944 (± 0.0029) | 0.7187 (± 0.0028) | 0.3180 (± 0.0051) | 0.6050 (± 0.0025) |
| | | 35 | 0.6663 (± 0.0022) | 0.7063 (± 0.0031) | <u>0.7432</u> (± 0.0032) | <u>0.3485</u> (± 0.0048) | 0.6081 (± 0.0021) |
| | | 150 | <u>0.6664</u> (± 0.0010) | <u>0.7065</u> (± 0.0017) | 0.7367 (± 0.0019) | 0.3479 (± 0.0021) | <u>0.6096</u> (± 0.0015) |
| | | 350 | 0.6534 (± 0.0017) | 0.7020 (± 0.0023) | 0.7217 (± 0.0022) | 0.3200 (± 0.0036) | 0.5961 (± 0.0020) |
| | | 650 | 0.6174 (± 0.0012) | 0.6644 (± 0.0020) | 0.6726 (± 0.0015) | 0.2424 (± 0.0026) | 0.5628 (± 0.0012) |
| | Half | 8 | 0.6480 (± 0.0021) | 0.6857 (± 0.0028) | 0.6986 (± 0.0028) | 0.2999 (± 0.0043) | 0.6169 (± 0.0023) |
| | | 35 | 0.6650 (± 0.0023) | 0.7086 (± 0.0023) | 0.7488 (± 0.0024) | 0.3519 (± 0.0047) | 0.5947 (± 0.0031) |
| | | 150 | 0.6701 (± 0.0016) | 0.7167 (± 0.0016) | 0.7539 (± 0.0014) | 0.3568 (± 0.0035) | 0.6116 (± 0.0017) |
| | | 350 | <u>0.6773</u> (± 0.0021) | <u>0.7316</u> (± 0.0017) | <u>0.7653</u> (± 0.0018) | <u>0.3762</u> (± 0.0044) | <u>0.6126</u> (± 0.0028) |
| | | 650 | 0.6429 (± 0.0021) | 0.6919 (± 0.0020) | 0.7164 (± 0.0017) | 0.2920 (± 0.0046) | 0.6022 (± 0.0016) |
| | Full | 8 | 0.6573 (± 0.0007) | 0.6994 (± 0.0024) | 0.7414 (± 0.0021) | 0.3323 (± 0.0019) | 0.5914 (± 0.0012) |
| | | 35 | 0.6832 (± 0.0009) | 0.7239 (± 0.0008) | 0.7695 (± 0.0010) | 0.3998 (± 0.0021) | 0.6041 (± 0.0017) |
| | | 150 | 0.6696 (± 0.0023) | 0.7200 (± 0.0027) | 0.7616 (± 0.0021) | 0.3519 (± 0.0052) | 0.6187 (± 0.0018) |
| | | 350 | <u>0.7046</u> (± 0.0017) | <u>0.7618</u> (± 0.0018) | <u>0.8056</u> (± 0.0014) | <u>0.4399</u> (± 0.0037) | <u>0.6381</u> (± 0.0021) |
| | | 650 | 0.6917 (± 0.0020) | 0.7438 (± 0.0020) | 0.7868 (± 0.0017) | 0.4107 (± 0.0038) | 0.6242 (± 0.0030) |

**Figure S3.**
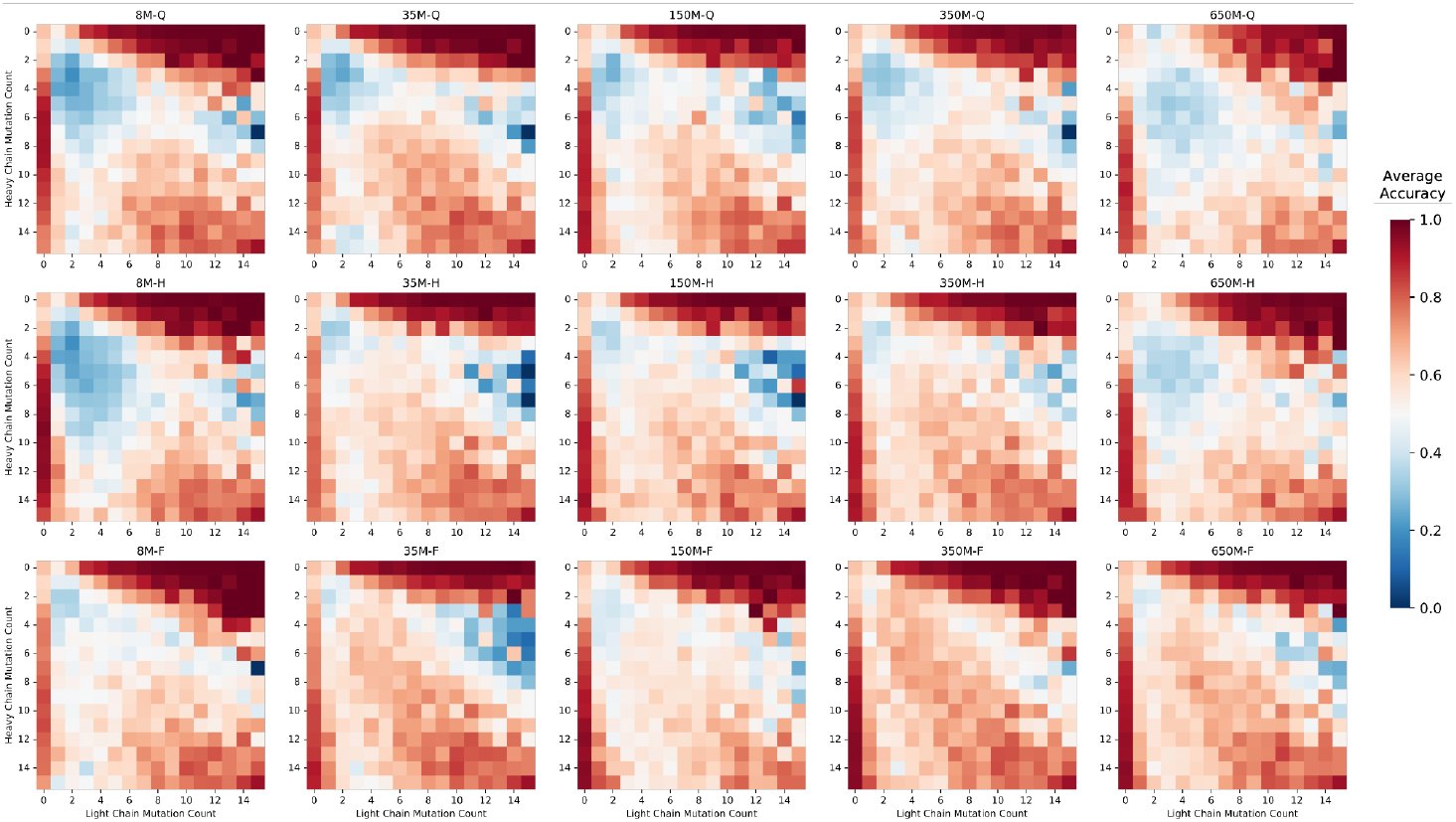
Classification accuracy across all model sizes and data scales for pair classification. Heatmaps show average classification accuracy for all models for different combinations of chain-specific mutation counts. Darker red values indicate higher accuracy, while lighter blue values indicate lower accuracy.

